# OGRE: Overlap Graph-based metagenomic Read clustEring

**DOI:** 10.1101/511014

**Authors:** Marleen Balvert, Tina Hauptfeld, Alexander Schönhuth, Bas E. Dutilh

**Author notes:** Shared last authors (Alexander Schönhuth, Bas E. Dutilh).

## Abstract

The microbes that live in an environment can be identified from the genomic material that is present, also referred to as the metagenome. Using Next Generation Sequencing techniques this genomic material can be obtained from the environment, resulting in a large set of sequencing reads. A proper assembly of these reads into contigs or even full genomes allows one to identify the microbial species and strains that live in the environment. Assembling a metagenome is a challenging task and can benefit from clustering the reads into species-specific bins prior to assembly. In this paper we propose OGRE, an Overlap-Graph based Read clustEring procedure for metagenomic read data. OGRE is the only method that can successfully cluster reads in species-specific bins for large metagenomic datasets without running into computation time-or memory issues.

## 1 Introduction

Metagenomics aims at identifying and characterizing the micro organisms that live in an environment by analyzing their combined genomic material. Next generation sequencing (NGS) technologies allow for the sequencing of genomic material at a relatively low cost, resulting in large amounts of short-read data. Sequencing a metagenome gives a large set of reads that originate from the genomes of a variety of species that live in the studied environment.

One of the most important general goals is to assemble the reads of the metagenome, that is to reconstruct the individual genomes that contribute to the metagenome. In doing so, one currently usually operates at the species level of taxonomy; that is, existing methods usually aim at reconstructing one genome for each of the species that contribute to the metagenome.

However, bacteria can already differ substantially at the strain level: consider a pathogen of which some strains are resistant to antibiotic treatment while others are not. To distinguish between resistant and non-resistant strains, and associate differences with (often clinically relevant) effects, knowing the differences at the genomic level is necessary. It is therefore desirable in metagenomics to assemble genomes not only at the species level, but to distinguish them at finer resolution and reconstruct genomes even at the strain level. Only this way, one is able to fathom the genetic diversity of a metagenome at sufficiently fine detail.

Recent work has pointed out, however, that assembling individual genomes from metagenomes at the strain level poses non-negligible methodical challenges. While some progress has been made already [Nurk et al., 2017, Baaijens et al., 2017], certain obstacles have been remaining. One most relevant such obstacle is that there are no methods that allow to cluster the reads from a metagenome into clusters that—at least with sufficiently high probability—only contain reads from identical or similar species. These clusters can then be picked up by assembly methods that are able to distinguish between individual genomes at the strain level. Thereby, it is important to note that such strain-level assembly methods usually encounter difficulties if the size of the input reads is too large. Clustering reads into species-specific groups scales down the problem in a meaningful way: a species cluster is often much smaller than the entire metagenome, while still collecting all reads that stem from identical species. Hence strain-specific genomes can be conveniently reconstructed by considering species-specific clusters in isolation.

Current methods, however, still rely on reducing the size of the input datasets by *assembling all reads into contigs first,* and only then, in *a second step, cluster reads into species-specific bins.* This procedure has a major flaw: since during the first step strain-specific cannot, but only consensus genome assembly methods can be used, differences at the strain level get lost. This, of course, hampers the entire procedure itself.

For reconstructing individual genomes from metagenomes at the strain level, novel methods are required. Following from the arguments from above, clustering reads into species-specific clusters *before employing assembly tools* would be highly desirable: after this step, strain-specific assembly tools, which exist but expect smaller-scale input, could conveniently pick up the resulting clusters and reconstruct the strains contributing to the species clusters.

Several reference-free binners for metagenomic short read datasets are available, all based on *k*-mer profiles. Short *k*-mers, where *k* depends on sequence length and abundance level [Wang et al., 2012], e.g. occur in the metagenomics read data at a frequency that is linearly proportional to their occurence in the genome they originate from, and the frequency increases with the abundance level of the species [Wu and Ye, 2011]. Abundancebin [Wu and Ye, 2011], TOSS [Tanaseichuk et al., 2012] and MBBC [Wang et al., 2015] use this property to derive the species’ abundances and bin the reads accordingly. The methods heavily rely on the assumption that no two species in the metagenome occur at similar abundance levels. MetaCluster 5.0 [Wang et al., 2012] bins the reads in three steps. First the reads from extremely low abundant (≤5x) species are filtered based on their *k*-mer frequencies. Second, MetaCluster 5.0 uses the observation that long *k*-mers are unique to a genome to bin reads originating from high-abundance species (>10x). The remainder of the reads (originating from low-abundance species) are then binned based on their *k*-mer profiles where *k* is intermediate (22 nt). Here, we apply these methods for read binning to the CAMI dataset ([Sczyrba et al., 2017], see Results). As none of the existing read binning methods performed well on the largest datasets (see Results), we developed a novel method that exploits Minimap2 [Li, 2017].

In this work we present OGRE, an Overlap Graph-based Read clustEring method. The intuition behind overlap-graph based clustering and assembly, and hence also our approach, is as follows: if one were to construct an overlap graph from reads that are sequenced from a single genome, then the graph consists of a single component when coverage over the entire genome is sufficiently high. On the other hand, when an overlap graph contains reads from different genomes (i.e. a metagenome) then no path exists in the overlap graph between two reads that originate from two different non-overlapping genomes. This is illustrated in Figure 1. An overlap graph is thus an intuitive tool for metagenomic read binning, where each connected component corresponds to a genome.

**Figure 1:**
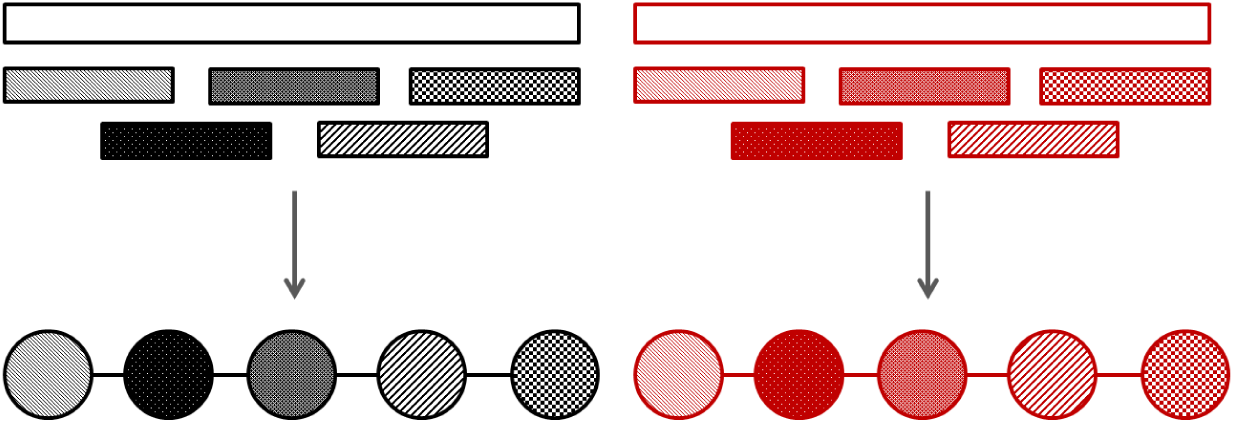
The two bars at the top represent genomes, the small bars map reads that are mapped back to these genomes. The color indicates the species. Reads that originate from the same genome form a single connected component in the overlap graph (provided that coverage is sufficiently high), while reads from different genomes are not connected in the overlap graph.

The idea of overlap graph-based clustering is similar to the idea behind MetaVelvet [Namiki et al., 2012]. This is an assembler designed for metagenomic datasets. A de Bruijn graph is constructed from the reads, which is decomposed into species-specific subgraphs by identifying chimeric nodes. Assembly is then performed on each of the de Bruijn sub graphs to obtain the genomes of the individual species. While this approach is similar to ours, there are some essential differences. First, while MetaVelvet employs a de Bruijn Graph, we make use of an overlap graph, which prevents the need for splitting reads into *k*-mers. Second, while MetaVelvet splits up the graph by searching for chimeric nodes, our graph is not connected to begin with and no splitting is needed: we only need to identify the connected components.

While the use of overlap graphs or de Bruijn graphs is common for assembly [Namiki et al., 2012, Nurk et al., 2017, Baaijens et al., 2017], no efficient implementation of an overlap graph-based clustering approach exists in the current literature. In this paper, we present OGRE, the first computationally feasible overlap graph-based read clustering approach for metagenomics data. OGRE employs Minimap2 [Li, 2017] which uses a clever heuristic approach for the efficient construction of an overlap graph. We show that OGRE is capable of clustering large metagenomic datasets without running into memory-or time issues, something which was not possible with existing clustering methods.

## 2 Methods

### 2.1 Clustering method

Despite the conceptual simplicity of overlap graph-based read clustering, its implementation is not straightforward: the algorithm needs to be able to efficiently construct an overlap graph from a large metagenomic dataset typically containing tens of millions of reads, and efficiently identify the clusters in the overlap graph. Here we first give a global overview of OGRE, further details are discussed in the following sections.

OGRE consists of three steps: (1) construct an overlap graph by identifying overlaps between reads, (2) from the list of overlaps select those that are expected to be an overlap between two reads from the same species, and (3) cluster the reads that are in the same connected component in the overlap graph (Figure 2). In the third step one may choose to impose a maximum cluster size to avoid unrealistically large clusters. For the overlap graph construction we use Minimap2 [Li, 2017], a recently developed tool that employs a heuristic approach to rapidly identify read overlaps. Minimap2 is the only tool that can identify the overlaps in a sufficiently short amount of time, and provides a list of overlaps between pairs of reads. For each overlap between two reads we compute a quality score to obtain an overlap graph with weighted edges, where higher weights correspond to a stronger overlap. Weights are in the interval [0,1], and edges (overlaps) with a weight below 0.5 are removed from the graph. We then cluster reads such that those pairs of reads with a high overlap score are in the same cluster. For this final step we use an efficient and parallel implementation of single linkage clustering. The overlap graph construction, overlap selection and clustering steps are further described in Sections 2.1.1, 2.1.2 and 2.1.3, respectively.

**Figure 2:**
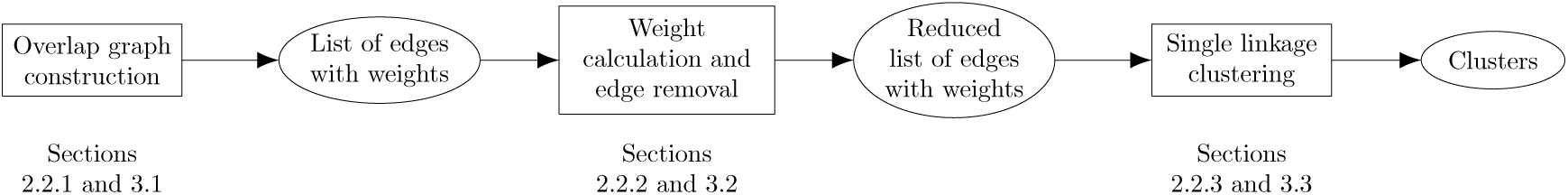
Workflow of the clustering method. Rectangles indicate steps in the workflow, ellipsoids indicate input/output to each of the steps. The sections in which the methods and results of each step are discussed are indicated below the steps.

#### 2.1.1 Overlap graph construction

Minimap2 outputs a list of pairs of overlapping reads with information corresponding to this overlap. It offers two possible output formats: a PAF file [Li, 2016] and a SAM file [Li et al., 2009]. We experienced some issues with the PAF output format, as it often reported only part of the overlap between two reads, which could then be mistaken for an overlap in the middle of a read and removed by our post-processing procedure. The SAM format does not suffer from this issue, as it outputs the CIGAR string from which the complete overlap range can be deduced. After running Minimap2 we discarded overlaps in the middle of reads as these make no sense in terms of clustering or assembly. This leaves us with a list of overlapping pairs of reads, hence a list of edges in the overlap graph. We chose settings for Minimap2 that allowed for identification of a large number of overlaps with a length of at least 60 bases (k=21, w=11, s=60, m=60, n=2, r=0, A=4, B=2, -end-bonus=100, see https://lh3.github.io/minimap2/minimap2.html for details).

The list of overlaps produced by Minimap2 contains overlaps where both reads originate from the same species as well as pairs where the reads come from different species. We would like to discard the latter. We aim to do so by predicting for each overlap whether the reads originate from the same species based on a measure of overlap strength. For this, we compute two metrics that could be relevant: the overlap length and a matching probability based on Phred scores (Appendix A, also used in Baaijens et al. [2017]). The predictive power of these metrics will be assessed in the results section.

Minimap2 produces a SAM file containing a large header and for each overlap the read IDs, CIGAR strings, mapping positions, segment sequence and Phred scores of the overlap. In our initial tests with a dataset (CAMI_low, see Section 3.1) containing 50,000,000 reads and hence 50,000,000^2^ pairs of reads, the output file exceeded 1TB - once the file reached a size of 1TB we stopped the process, so we don’t know how large it could get. However, for our purposes we only need a list of overlaps characterized by the read IDs, the overlap length and the Phred-based overlap quality score, which requires less space. To prevent the production of a large output file we split the set of reads into *n* equal subsets *s*_1_,…, *s_n_* and ran Minimap2 for every pair of subsets (*s_i_*, *s_j_*) ∈ {*s*_1_,…, *s_n_*} × {*s*_1_,…, *s_n_*}. As Minimap2 produces a slightly different output when running it on (*s_i_*, *s_j_*) compared to (*s_j_*, *s_i_*), i ≠ *j*, that is, it matters which subset is used as a reference and which one as a query, we ran both, resulting in *n*^2^ small output files. For each file we only kept the read pair IDs, CIGAR strings, read sequences and Phred scores and as such removed a lot of redundant information. We then computed the overlap length and the Phred-based matching probability from the CIGAR strings, the read sequences and Phred scores, after which the latter three were discarded.

Next, for each pair of subsets, we combined the two overlap files (one obtained with the first subset as a reference and the second as a query, and one obtained with the reverse) and removed all overlaps occurring twice. We now have *n* + (*n* — 1) + … + 2 + 1 = *n* * (*n* + 1)/2 overlap files. These were merged into a single overlap file that contained a list of overlaps characterized by the corresponding read IDs, overlap length and Phred-based overlap score. This file is the list of edges in our overlap graph.

#### 2.1.2 Filtering overlaps between reads from different species

As already mentioned we wish to give each edge a weight that indicates the strength of the overlap, and filter those overlaps for which the reads originate from different species from the full list of overlaps obtained. For this purpose we use a logistic regression, a machine learning method that predicts for each sample a binary class based on a sigmoid function applied to a linear transformation of sample characteristics:

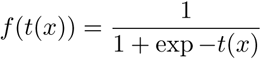

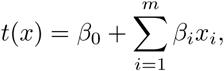

where *x* ∈ 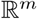 is the vector of sample characteristics. Note that *f*(*t*(*x*)) ∈ [0*and*1]. Samples with a value >0.5 are predicted to be in one class, the other samples in another. In our case, the classes are “same species” and “different species”, where *f*(*t*(*x*)) > 0.5 corresponds to “same species”, and the characteristics are the overlap length and the Phred based overlap score. The output value of the logistic regeression is used to obtain edge weights in the interval [0,1], and edges that have a weight below 0.5 were removed. Values for the parameters β_0_,…,β_*m*_are obtained by training the model on past data where the true class of each sample is known. We further discuss our approach to this in Section 3.6.

#### 2.1.3 Implementation of the single linkage clustering algorithm

The list of overlaps - constructed as described above - describes the overlap graph and thus forms the starting point of the clustering algorithm. Initially each read forms a bin by itself, and in every iteration two bins are merged. The single linkage algorithm chooses the two bins with the largest overlap score between them and merges them. The overlap score between two bins *A* and *B* is defined as the maximum overlap score over all pairs of read ends *r* = (*r*_1_*,r*_2_)*,r*_1_∈ *A*, *r*_2_∈ *B* (Figure 3). Sorting the overlaps according to decreasing overlap score provides the order in which bins should be merged. After sorting, the remainder of the merging algorithm is linear in the number of edges: for each line in the file, the two bins that contain the reads of that line are merged.

**Figure 3:**
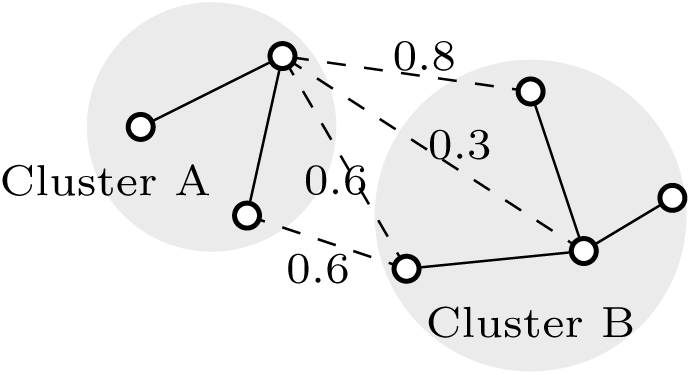
Example of the single linkage clustering. The solid lines denote edges between nodes in the same cluster, the dashed lines denote edges between nodes from different clusters with their overlap scores. The overlap score between clusters A and B is the maximum overlap score over the four connections between the clusters, which equals 0.8.

When we are looking for the connected components in a graph, all edges will be used, so the order in which bins are merged may seem irrelevant. However, a connected component in an overlap graph may contain multiple, possibly many, species when they share part of their genome sequence. This may lead to huge clusters containing reads from many genomes. In order to prevent this, we limit the cluster size: reads are clustered in order of decreasing overlap strength, and a cluster is considered “full” and removed from the clustering procedure when its size reaches the threshold. In this case the edge weights are relevant.

Although the above algorithm has a complexity of 𝒪(*n*log *n+n*) where *h* is the number of edges in the graph (the complexity of the merge sort algorithm that is used by the bash command “sort” plus the complexity of the single linkage algorithm), the size of the overlap graph makes executing it nontrivial: any serial clustering algorithm, even a very fast one, will have to go through all the edges that are in the list of overlaps and will thus take a very long time to process. In order to be able to use the methodology described above, we use several techniques to speed up the method.

##### Hash table

We store our clustering in a hash table where each key is a node (read) and its value is the ID of the cluster the read belongs to (see left part of Figure 4a). As noted before, initially each read forms a cluster on its own. When merging two (single-node) clusters we update the value of only one node to be equal to the ID of the other node instead of the cluster ID. This results in a cluster with two nodes that is stored as a chain of keys and values (Figure 4a). When merging two clusters that contain multiple nodes we update only the value of the node that points to the cluster ID, also referred to as the head of the cluster, in one of the clusters to be equal to the head node of the other cluster (Figure 4b). As a result, only a single value needs to be updated whenever two (potentially large) clusters are merged.

**Figure 4:**
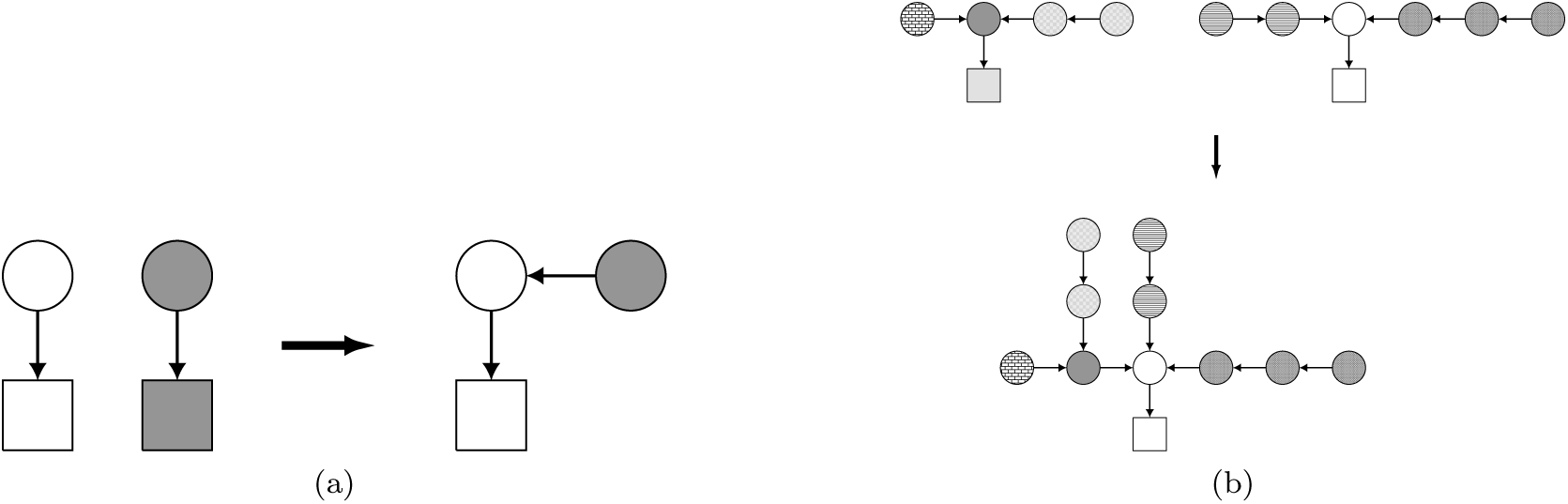
Merging two clusters of (a) a single node each and (b) multiple nodes by changing the value of one node to be equal to the ID of the other node. Circles indicate reads, squares indicate cluster IDs. An arrow points from a key to its value in the hash table. An arrow from a circle to a square indicates that this read is in the cluster corresponding to the square. An arrow from one circle to the next indicates that the read from which the arrow departs is in the same cluster as the read that the arrow points to. Fill patterns allow for identification of nodes between the two steps.

##### Updating the cluster with the shortest maximum chain

Suppose that we have arrived at an iteration where the algorithm considers the overlap between reads *r*_1_ and *r*_2_. Then the clusters containing reads *r*_1_ and *r*_2_, denoted by *C*_1_ and C_2_, respectively, need to be merged. We need to identify the heads of *C*_1_ and *C*_2_ by traversing the path through the hash table starting at reads *r*_1_ and *r*_2_ towards the respective cluster heads. The number of operations for this procedure equals the length of the path from *r*_1_ to the head of cluster *C*_1_ plus the length of the path from *r*_2_ to the head of cluster *C*_2_. For example, consider the cluster in the lower part of Figure 4b, and suppose that the right-most node in this cluster is *r*_1_. In order to find the cluster ID (the square) one needs to traverse three nodes, hence four operations are required to arrive at the cluster ID. It is thus desirable to keep the length of the longest chain in each cluster as short as possible. When merging two clusters we therefore redirect the head of the cluster with the shortest maximum chain within the cluster to the head of the other cluster. As a result, the maximum chain within a cluster of size *n* will never exceed 1 + [*n/*2] (see Appendix B).

##### Parallellization of the single linkage clustering algorithm

Although the single linkage clustering algorithm is very fast and simple, the list of overlaps contains from tens of millions up to billions of lines and going through these one by one could take up to weeks or months for a dataset of the size that is often encountered in metagenomics. We thus need to parallelize the approach. As the merging procedure is sequential by nature, it is difficult to parallelize the merging itself. However, note that after several iterations many lines in the list of overlaps will become redundant: they are overlaps between pairs of reads that either are already in the same cluster, or merging their clusters yields a too large new cluster. This observation forms the basis for our parallelization method. In each iteration, each of the parallel processes receives a block of lines (e.g. 1,000,000 lines) retrieved from the position in the list of overlaps to where the algorithm got so far. For each of these lines the process checks (1) whether the two reads are in a different cluster and (2) whether merging their clusters leads to a cluster size that does not exceed the threshold. If both conditions are met, the line is written to a new list. After all processes have finished, the clustering algorithm only needs to consider those pairs of reads that were written to the lists by the separate processes. The number of lines processed in each iteration is thus 1,000,000 × the number of processes. After a few iterations, the number of overlaps that makes it to the filtered list is only a fraction of the total number of pairs of reads processed by the individual processes. An overview of this approach is given in Figure 5.

**Figure 5:**
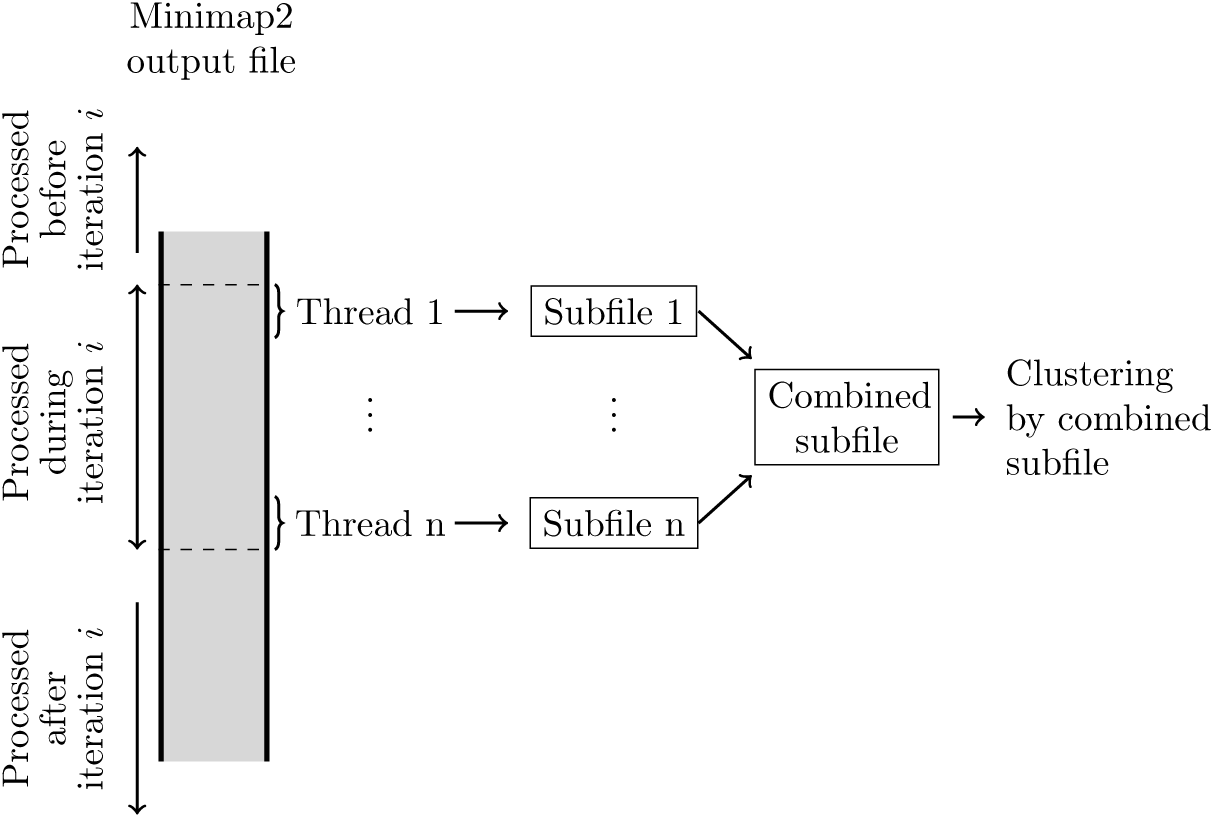
Parallelization of the single linkage clustering: processes as executed in a single iteration *i*.

## 3 Results

### 3.1 Data

We test OGRE on the datasets provided by the CAMI challenge [Sczyrba et al., 2017]. Six datasets are provided, containing simulated short Illumina reads from a mixture of strains. In all but one dataset these strains can be grouped into species (for one dataset there is only a single strain per species present). The datasets differ in complexity. Summary statistics are given in Table 1, further details can be found in Sczyrba et al. [2017].

**Table 1:** Characteristics of six CAMI datasets. ^*^Four out of the six CAMI datasets contain multiple samples. For these we only used samples S001. ^**^This is a relative complexity as indicated by Sczyrba et al. [2017], see their paper for further details.

| Dataset | Sample* | No. reads | Read length (nt) | No. species | No. strains | Complexity** |
| --- | --- | --- | --- | --- | --- | --- |
| CAMI_low | - | 49,898,179 | 150 | 27 | 60 | Low |
| CAMI_medium | S001 | 66,489,042 | 150 | 91 | 232 | Medium |
| CAMI_high | S001 | 49,901,367 | 150 | 376 | 1074 | High |
| toy_low | - | 72,855,674 | 100 | 30 | 30 | Low |
| toy_medium | S001 | 77,155,802 | 100 | 199 | 225 | Medium |
| toy_high | S001 | 74,016,648 | 100 | 375 | 450 | High |

### 3.2 Clustering CAMI_low with available read clustering methods gives memory and time issues

We have attempted to perform read clustering of the CAMI_low dataset (Table 1) using Abundancebin [Wu and Ye, 2011], TOSS, [Tanaseichuk et al., 2012], MetaCluster 5.0 [Wang et al., 2012] and MBBC [Wang et al., 2015]. These methods were unable to finish within reasonable time. The methods were applied to subsets of the data. MBBC spent over a month on a subset containing 5% of the species before we stopped the process. Running Abundancebin on a subset containing 5% of the species was possible, but after running it on 20% of the species for three weeks we stopped the process. As TOSS depends on Abundancebin, we have not tried running it separately. MetaCluster could finish running on 20% of the data, but was not able to handle the full dataset.

### 3.3 Runtimes for the clustering approach are acceptable for all but one dataset

Runtimes for the overlap graph construction and the clustering step are presented in Table 2. Note that the toy_low dataset (Table 1) is missing: our method could not be run on this dataset within reasonable time. This is due to the large number of overlaps that Minimap identifies, which makes it impossible to obtain the full overlap graph without running into memory and time issues. Therefore, no further results were obtained for this dataset. For the remaining five datasets however our parallelization approach makes overlap graph-based clustering feasible on a multi-core system. Our results were obtained on a system with 24 CPUs, and were all finished within a week.

**Table 2:** Run time in CPU hours of the algorithm.

| Dataset | Run time (CPU hours) |  |  |
| --- | --- | --- | --- |
|  | Overlap graph construction | Clustering | Total |
| CAMI_low | 2118 | 145 | 2263 |
| CAMI_medium | 1853 | 367 | 2220 |
| CAMI_high | 291 | 250 | 541 |
| toy_medium | 1762 | 291 | 2053 |
| toy_high | 648 | 24 | 672 |

### 3.4 Overlap graph construction can be done without memory issues

The main issue with constructing an overlap graph by directly running Minimap2 on the CAMI datasets is the size of the resulting file, as this yields an excessively large output file. The algorithm described in Section 2.1.1 was developed to overcome this issue, hence the main performance indicator for this step is the size of the resulting overlap files. As Table 3 shows, the final overlap file has an acceptable size.

**Table 3:** Number of edges in the overlap graph and size of the output file.

| Dataset | Number of edges<br>($\times 10^9$) | Output file size<br>(GiB) |
| --- | --- | --- |
| CAMI_low | 8.10 | 310 |
| CAMI_medium | 7.04 | 360 |
| CAMI_high | 0.36 | 19 |
| toy_medium | 2.85 | 137 |
| toy_high | 2.33 | 108 |

### 3.5 The prediction step removes many different species overlaps and keeps most same species overlaps

For a given overlap we aim to predict whether the corresponding reads originate from the same genome based on the overlap length and a Phred-based matching probability.

First we compare the distributions of the overlap length and the Phred-based matching probability for same-species pairs of reads versus different-species pairs of reads. We randomly selected 2,000,000 overlaps from each of the datasets: 1,000,000 for which the reads are from the same species and 1,000,000 for which the reads originate from different species. The distributions for overlap length and Phred-based matching probability were plotted for same-species overlaps (blue) and different-species overlaps (red) separately (Figure 6 for CAMI_low and Figures 9 up to 12 in Appendix C for the remaining datasets). The histograms show that the Phred-based probability score can be a valuable predictor for whether the overlap corresponds to reads originating from the same species, while the relation between overlap length and read origin seems to be less indicative.

**Figure 6:**
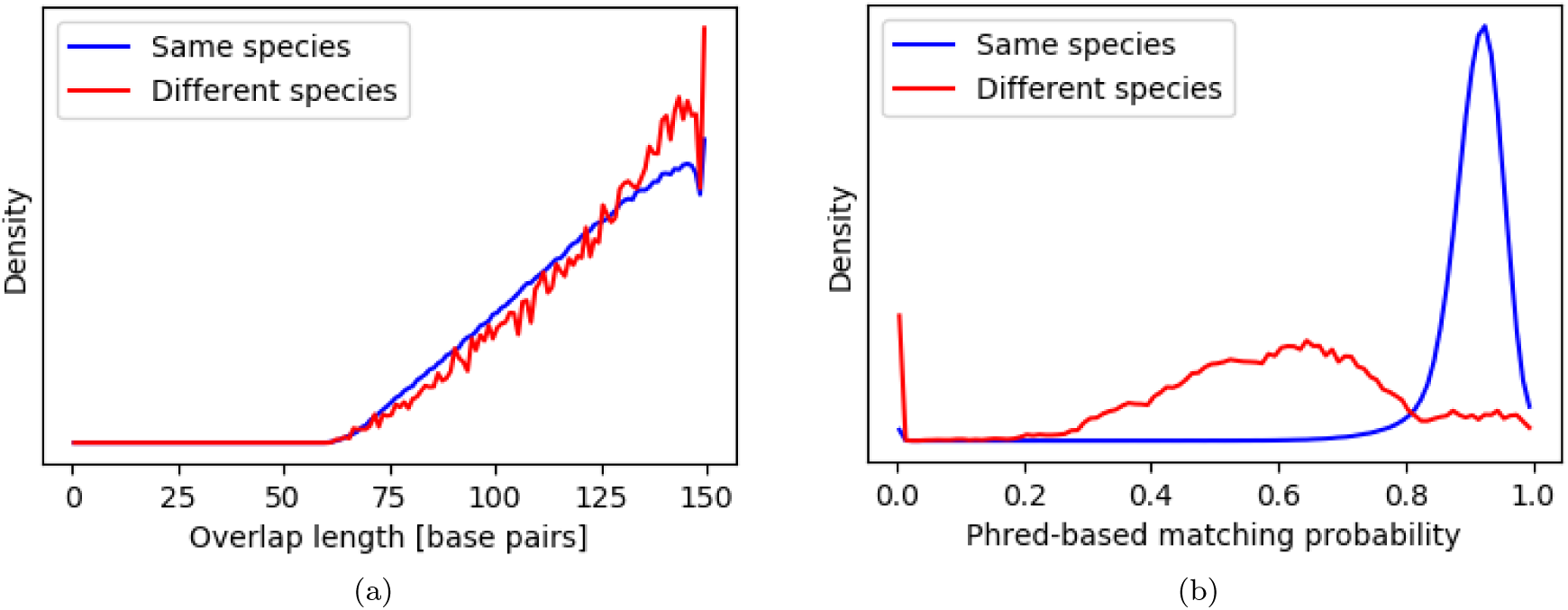
Distribution of (a) the overlap length and (b) the Phred-based matching probability for 1,000,000 overlaps between reads from the same species and 1,000,000 overlaps between reads from different species randomly selected from CAMI_low

Recall that we aim to discard overlaps between reads from different species, while retaining as many same species overlaps as possible using a logistic regression. The model needs to be trained on synthetic data where for each read it is known from which species it originates. The CAMI datasebase is highly suitable for this purpose. We trained our models based on all overlaps from four out of the five datasets. The performance of the trained models was tested using the fifth dataset, which has a different complexity than the other datasets and was withheld completely up to this point.

Table 4 shows train and test accuracies for each of the five datasets. The dataset in the first column is the dataset for which the reads are to be clustered (the test dataset). The training dataset is constructed by randomly selecting 10,000 overlaps between reads from the same species and 10,000 overlaps between reads from different species from each of the four datasets other than the test dataset, and combining those into a single training dataset. Note that there is a big gap between the training and the test accuracy, which is due to a combination of two factors. The test dataset (containing the complete overlap graph) is highly unbalanced: over 99% of the edges correspond to an overlap between reads from the same species. This, combined with the model being far better at recognizing same species overlaps than different species overlaps, yields a higher accuracy for the test data than for the training data.

**Table 4:** Training and test accuracies. The training data for the dataset in column 1 is created by selecting 10,000 same species overlaps and 10,000 different species overlaps from each of the overlap graphs of the five datasets other than the test dataset. Test accuracy is obtained by applying the trained model to the overlap graph of the test dataset.

| Test dataset | Train accuracy | Test accuracy |
| --- | --- | --- |
| CAMI_low | 0.715 | 0.801 |
| CAMI_medium | 0.775 | 0.923 |
| CAMI_high | 0.765 | 0.891 |
| toy_medium | 0.758 | 0.925 |
| toy_high | 0.772 | 0.934 |

The goal of this step in the clustering procedure is to discard as many overlaps between reads from different species as possible - a single such overlap leads to merging two clusters with reads from different species - while keeping most of the overlaps between reads from the same species. We therefore look at the fraction of same species-and different species overlaps that were discarded by the logistic regression classifier (Table 5). The results show that while for most datasets over 90% of the same species overlaps were kept, around half of the overlaps between reads from different species were discarded, and for CAMI_low this was even over 90%.

**Table 5:** Fraction of overlaps discarded by the logistic regression classifier.

| Test dataset | Same species overlaps | Different species overlaps |
| --- | --- | --- |
| CAMI_low | 0.199 | 0.935 |
| CAMI_medium | 0.077 | 0.446 |
| CAMI_high | 0.109 | 0.561 |
| toy_medium | 0.075 | 0.598 |
| toy_high | 0.066 | 0.483 |

### 3.6 Clustering gives sensible results for all datasets but one

For each dataset the clustering was performed with a maximum cluster size of 3300, 17000 and 33000 reads (corresponding to at most 1 million, 5 million and 10 million base pairs, respectively), as well as with unlimited cluster size. All clusters of fewer than 20 reads were discarded, and these reads were considered as not clustered.

Detailed results of CAMI_low are shown here. The results of the CAMI_medium, CAMI_high and toy_medium datasets are comparable to those of CAMI_low, and are thus not separately discussed. Their results can be found in Appendices D up to F. The results of toy_high will be discussed here, as they are not comparable to those of the other datasets.

In a perfect clustering each read is clustered, all reads from the same species are in the same cluster and each cluster contains reads from only one species. We will therefore assess the number of clustered reads, the number of clusters per species and the number of species per cluster.

First we plotted the fraction of the reads that was clustered versus coverage for each species in a dataset, see Figures 7 (CAMI_low and toy_high) and 13 in Appendix D (remaining datasets), where each dot represents a species. Results for the four maximum cluster sizes are shown in red (3300 reads), blue (17000 reads), green (33000 reads) and black (unlimited). The figures show that the fraction of reads that was clustered is generally high for species with a high coverage. Furthermore, as the maximum cluster size increases, the number of clustered reads increases as well. The exception is toy_high, where hardly any reads were clustered at all.

**Figure 7:**
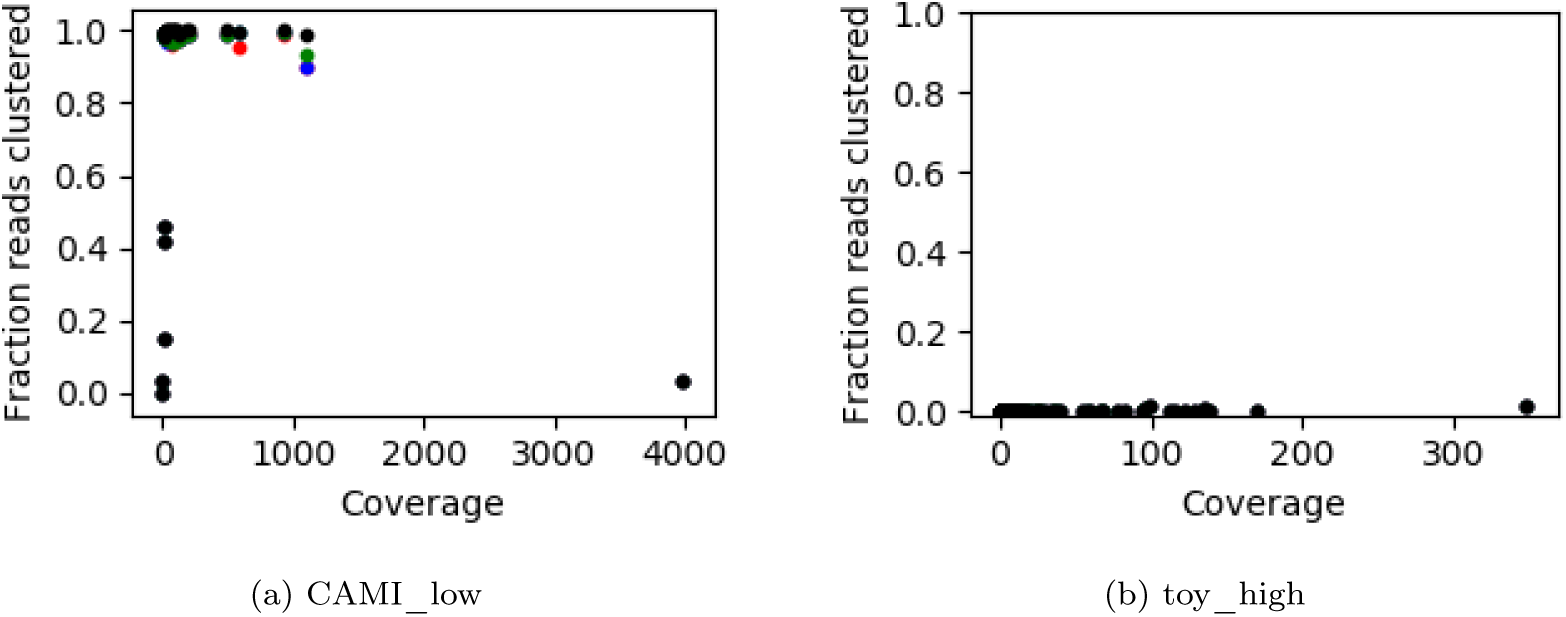
Fraction of the reads that was clustered versus coverage for (a) CAMI_low and (b) toy_high. Each dot represents a species in the dataset. Results are presented for the four maximum allowed cluster sizes: 3300 reads (red), 17000 reads (blue), 33000 reads (green) and no limit (black).

Tables 6 (CAMI_low) and 7 (toy high_compexity) show the distributions of the number of species in a cluster for the four maximum cluster sizes. The same tables for the remaining datasets can be found in Appendix E. The tables show that as the maximum allowed cluster size increases, the maximum number of species in a cluster increases as well. On the other hand, the total number of clusters decreases.

**Table 6:** Number of clusters containing the number of species indicated in the first column for CAMI low for the four maximum cluster sizes.

|  |  | Max cluster size (reads) |  |  |  |
| --- | --- | --- | --- | --- | --- |
|  |  | 3300 | 17000 | 33000 | Unlimited |
| No. species in cluster | 1 | 25466 | 12314 | 10224 | 4235 |
|  | 2 | 25 | 26 | 22 | 4 |
|  | 3 | 5 | 2 | 3 | 0 |
|  | 4 | 1 | 0 | 0 | 0 |
|  | 5 | 1 | 0 | 0 | 0 |
|  | 7 | 0 | 1 | 0 | 0 |
|  | 11 | 0 | 0 | 1 | 0 |
|  | 15 | 0 | 0 | 0 | 1 |
| Total |  | 24498 | 12343 | 10250 | 42340 |

**Table 7:**
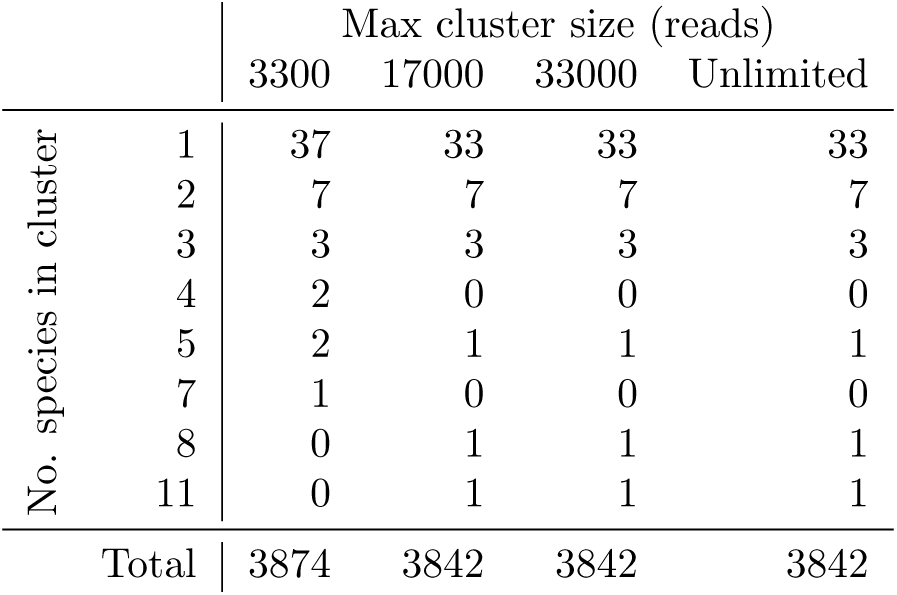
Number of clusters containing the number of species indicated in the first column for toy_high for the four maximum cluster sizes.

|  |  | Max cluster size (reads) |  |  |  |
| --- | --- | --- | --- | --- | --- |
|  |  | 3300 | 17000 | 33000 | Unlimited |
| No. species in cluster | 1 | 37 | 33 | 33 | 33 |
|  | 2 | 7 | 7 | 7 | 7 |
|  | 3 | 3 | 3 | 3 | 3 |
|  | 4 | 2 | 0 | 0 | 0 |
|  | 5 | 2 | 1 | 1 | 1 |
|  | 7 | 1 | 0 | 0 | 0 |
|  | 8 | 0 | 1 | 1 | 1 |
|  | 11 | 0 | 1 | 1 | 1 |
| Total |  | 3874 | 3842 | 3842 | 3842 |

The number of clusters per species versus coverage is shown in Figure 8 for CAMI_low and toy_high, and in Figure 14 in Appendix F for the four remaining datasets. As before, each dot represents a species, and the red, blue, and green dots correspond to a maximum allowed cluster size of 3300, 17000 and 33000 reads, while black dots correspond to an unlimited cluster size. As can be seen in the figure, the reads from species with a low coverage are spread over a larger number of clusters and hence are less well clustered.

**Figure 8:**
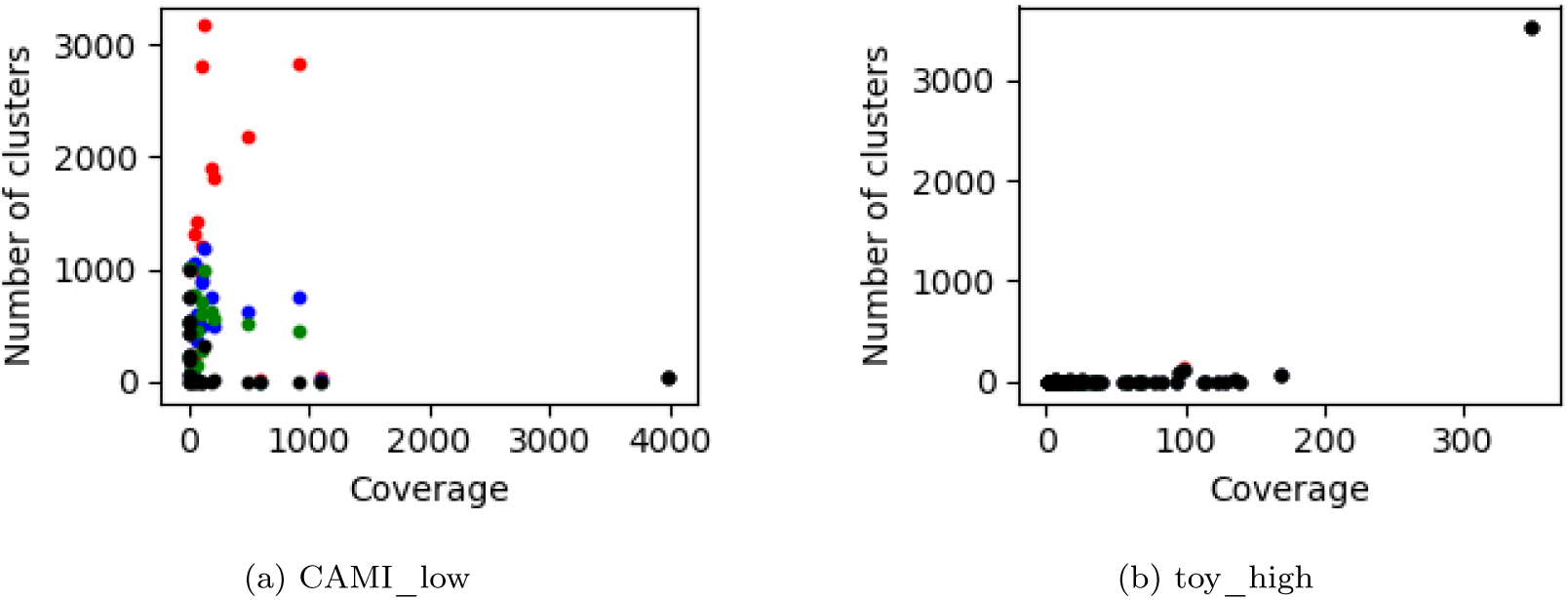
The number of clusters that contain at least one read from a species versus coverage for (a) CAMI_low and (b) toy_high. Each dot represents a species in the dataset. Results are presented for the four maximum allowed cluster sizes: 3300 reads (red), 17000 reads (blue), 33000 reads (green) and no limit (black).

### 3.7 Weak clustering for toy_high is caused by complexity of the dataset

The toy_high dataset is the only dataset for which a weak clustering was obtained. In order to see what caused this, we assembled all datasets using MetaSpades [Nurk et al., 2017] and investigated the quality of the assembly. The results in Table 3 show that the assembly of toy_high is of much lower quality in terms of mean and maximum contig length, N50 and N80 than for the four other datasets. The complexity of the dataset causes both MetaSpades and our approach to have difficulties.

**Table 8:**
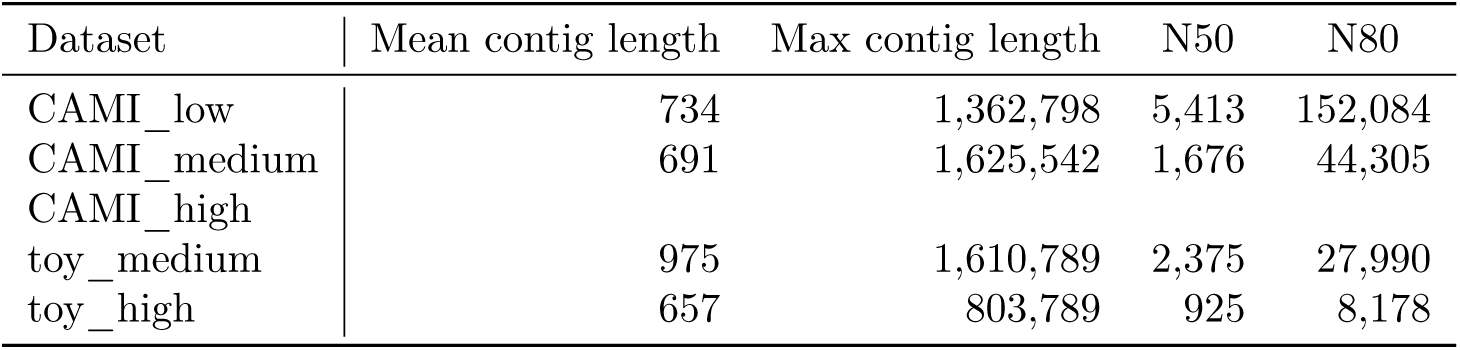
Maximum and mean contig length, N50 and N80 for the assembly performed with Metaspades.

| Dataset | Mean contig length | Max contig length | N50 | N80 |
| --- | --- | --- | --- | --- |
| CAMI_low | 734 | 1,362,798 | 5,413 | 152,084 |
| CAMI_medium | 691 | 1,625,542 | 1,676 | 44,305 |
| CAMI_high |  |  |  |  |
| toy_medium | 975 | 1,610,789 | 2,375 | 27,990 |
| toy_high | 657 | 803,789 | 925 | 8,178 |

## 4 Discussion

This paper presents OGRE, an overlap graph-based read clustering approach for clustering the reads in a large metagenomic dataset. For this several challenges were resolved. First, the strongest (and probably only suitable) overlap graph construction tool that is available, Minimap2, results in an unacceptably large overlap file for our test cases. This issue was overcome by applying Minimap2 on parts of the data, sytematically removing redundant information from the separate overlap files, and systematically merging the resulting files. Second, while the single linkage algorithm is a highly efficient approach for clustering the reads from the overlap graph, it is sequential in nature and applying it to the long edge list we obtained with Minimap2 results in unacceptable computation times. The parallelization approach presented here allowed us to overcome this issue. Third, the overlaps obtained with Minimap2 contain a large number of overlaps between reads from a different species. About half of these overlaps could be filtered out using a logistic regression on the overlap length and a Phred-based matching probability.

The presented approach could not provide a clustering for toy_low due to the large number of overlaps that Minimap2 identifies for this dataset. One can tune the parameters of Minimap2 such that it stores fewer overlaps, e.g. by increasing the minimum overlap length or the threshold on mismatch acceptance. As a result, the fraction of same species overlaps may increase which is expected to strengthen the clustering. On the other hand, no proper clustering was obtained for toy_high. This may be overcome by loosening the restrictions on Minimap2, resulting in a larger overlap file. Setting the parameters for Minimap2 is thus a dataset-dependent task.

In practice, there are two difficulties with the concept of overlap graph-based read clustering. First, a connected component in the overlap graph may contain reads from multiple genomes that share part of their sequence, which may have occured through e.g. horizontal gene transfer or domain conservation. In our experiments this has resulted in some clusters containing reads from multiple species. Second, reads from a single genome may cluster into multiple separate connected components in the overlap graph when coverage is low. This is consistent with our observations: while the clustering approach works well for species with a relatively high coverage, many reads from species with low coverage (below approximately 15x) remain unclustered. The issue may be overcome by applying a *k*-mer based read clustering method to the remaining reads. For example, MetaCluster 5 [Wang et al., 2012] provides an approach to identify some the species with extremely low abundance. This is left for future research.

A species-specific read clustering approach for metagenomics such as OGRE paves the way for faster assembly [Namiki et al., 2012] as this allows for genome assembly for species-specific bags of reads. We foresee great value in combining our read clustering approach with an assembly tool such as SPAdes [Bankevich et al., 2012] or Savage [Baaijens et al., 2017].

## 5 Conclusion

This paper presents OGRE, an overlap-graph based read clustering approach. While this is a computationally intensive task, we developed a parallelized approach such that an overlap graph-based method becomes feasible even for realistic large metagenomic datasets. This makes it the only de novo read clustering method that can handle datasets of such size. While OGRE was capable of providing a good read clustering for four out of the six test datasets, no proper clustering was obtained for two datasets. This indicates the delicacy of setting the Minimap2 parameters, for which no generally applicable values can be chosen. Overall, we conclude that the presented method is a strong first step towards further development of overlap graph-based read clustering.

## Acknowledgements

### Funding

MB is supported by the Netherlands Organization for Scientific Research (NWO Vidi grants 639.072.309 and 864.14.004. AS is supported by the Netherlands Organization for Scientific Research (NWO Vidi grant 639.072.309. BED is supported by the Netherlands Organization for Scientific Research (NWO) Vidi grant 864.14.004.

### A Calculation of the Phred-based matching probability

Suppose we have an overlap for which we observe the two read sequences *s*_1_*, s*_2_∈ {*A,C, G, T*}^*n*^ with corresponding Phred scores *p*_1_*,p*_2_∈ {ascii}^*n*^. While *s*_1_ and *s*_2_ denote the *observed* read sequences, we denote the true (unknown) read sequences by σ_1_and σ_2_. The probability that *s*_1_ and *s*_2_ have the same base pair in the *i^th^* position can be calculated as follows:

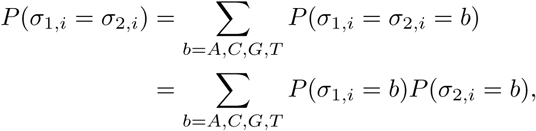

where

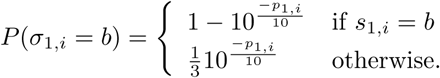

From this we compute the Phred-based matching probability, which is defined as the probability that two bases are identical averaged over the full overlap:

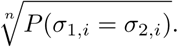

### B Maximum chain length in a cluster

#### Proposition

*The maximum chain within a cluster of size n will never exceed* 1 + [n/2]

##### Proof

The validity of the proposition can be shown by induction. First, note that this holds for *n* =1: if a cluster contains one node, than that one node will directly point to the cluster ID and the chain has length 1. Now assume that the statement holds for clusters of size at most *n* — 1, that is, the longest chain in a cluster of size *n* — 1 is of length 1 + [(*n* — 1)/2]. Consider a cluster of size *n*, which we denote as cluster *A*. This cluster was formed by merging two clusters, say clusters *B* and *C* with *n_B_* and *n_C_* nodes, *n_B_*≤ *n*/2, *n_C_*≥ *n*/2. From these two clusters, we redirected the pointer of the head of the cluster with the shortest maximum chain, let’s say that this chain has length *l*. The maximum chain in the new cluster has length *l* + 1 by construction. By assumption, the length of the maximum chain in cluster *B* does not exceed 1 + [*n_B_* /2]. For *n* = 2 we have *n_B_ = n_C_ =* 1, while for *n* = 3 we have *n_B_* = 1 and *n_C_ =* 2. In both cases this gives 1 + [*nB*/2] = 1 = *n*/2. When n ≥ 4 we can write:

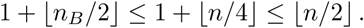

Hence for any *n* > 1 we have *l*≤ [*n*/2] and the maximum chain in the new cluster has a length of at most 1 + [*n*/2].

### C Distributions of overlap metrics

**Figure 9:**
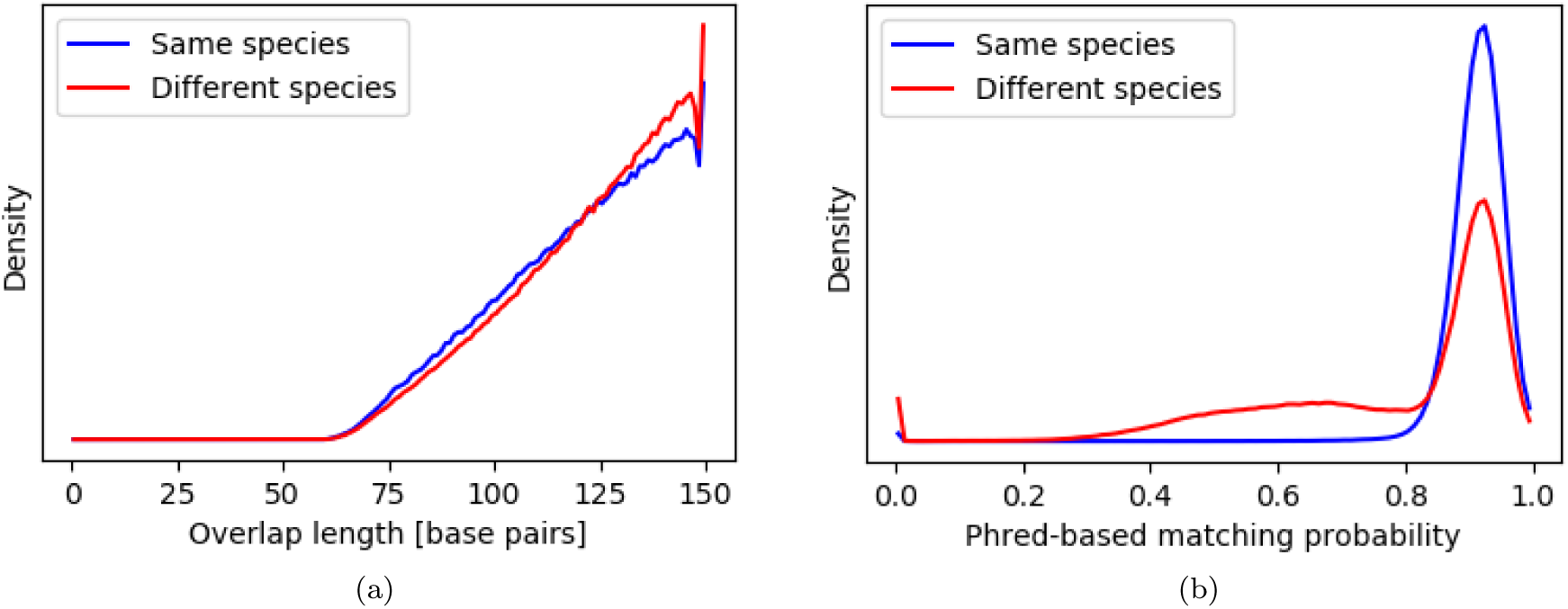
Distribution of (a) the overlap length and (b) the Phred-based matching probability for 10,000 overlaps between reads from the same species and 10,000 overlaps between reads from different species selected from CAMI_medium.

**Figure 10:**
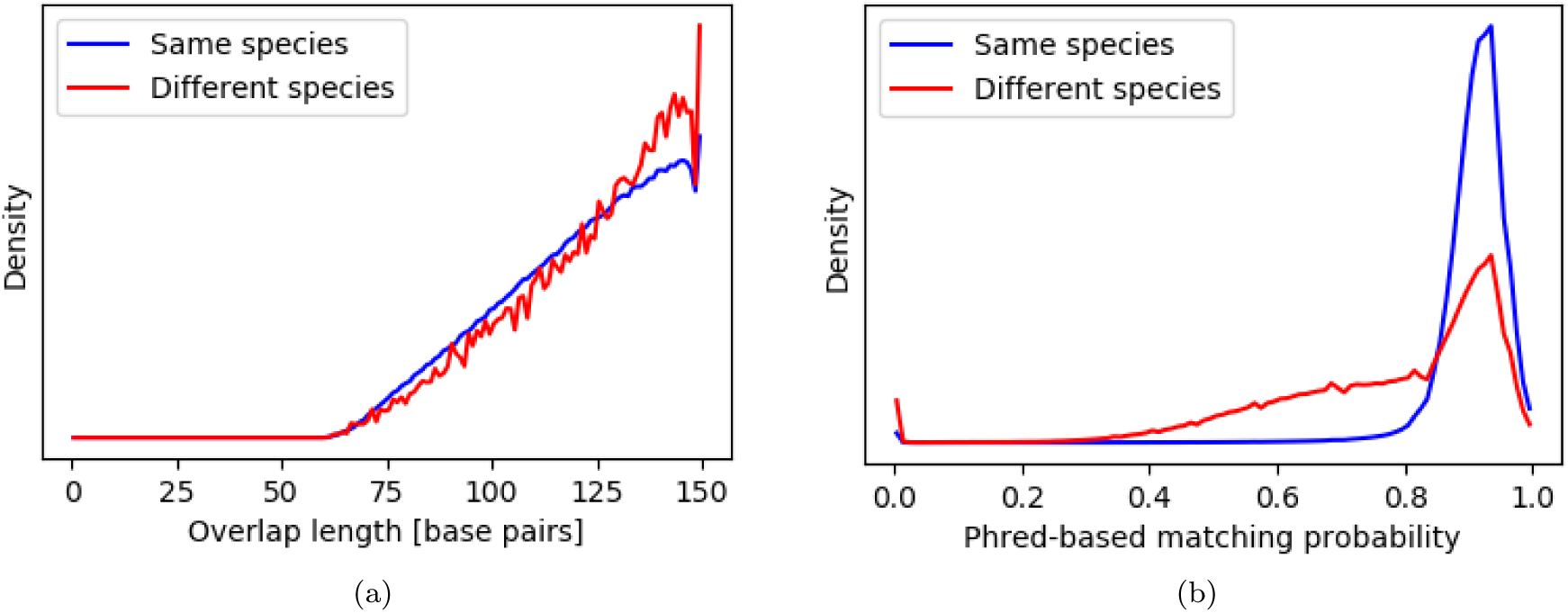
Distribution of (a) the overlap length and (b) the Phred-based matching probability for 10,000 overlaps between reads from the same species and 10,000 overlaps between reads from different species randomly selected from CAMI_high.

**Figure 11:**
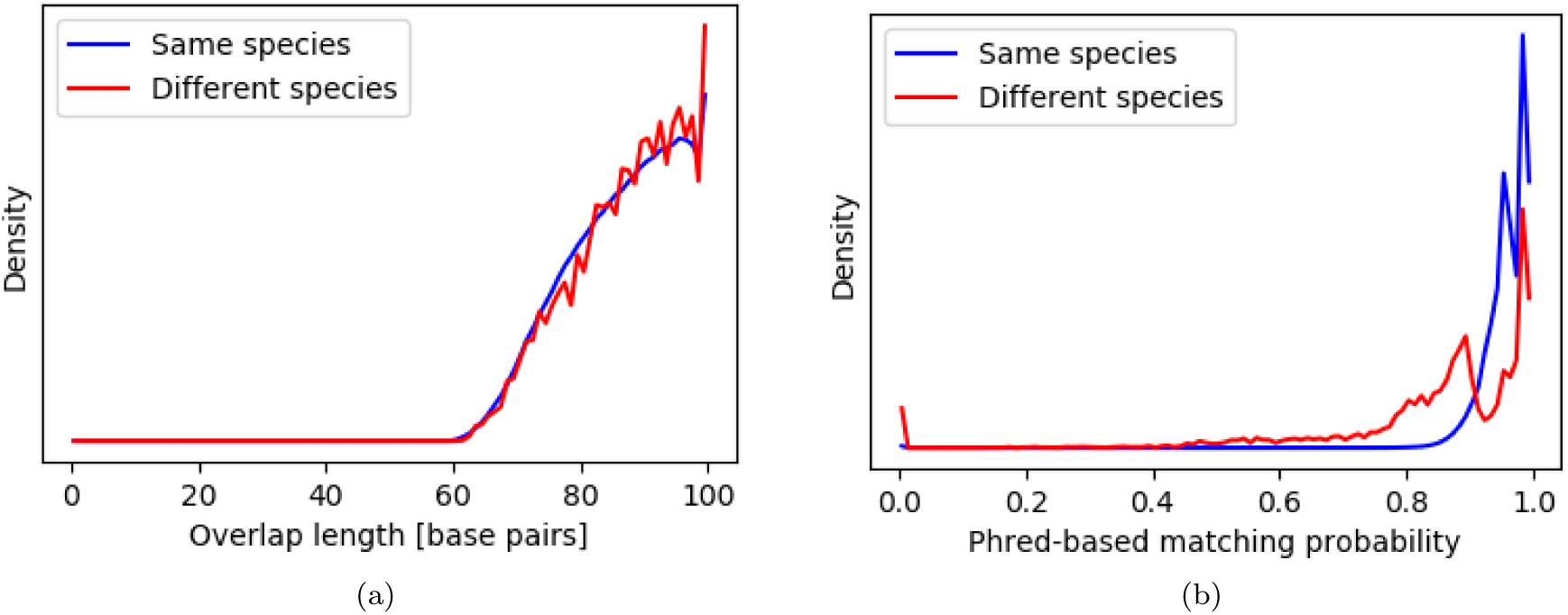
Distribution of (a) the overlap length and (b) the Phred-based matching probability for 10,000 overlaps between reads from the same species and 10,000 overlaps between reads from different species randomly selected from toy_medium.

**Figure 12:**
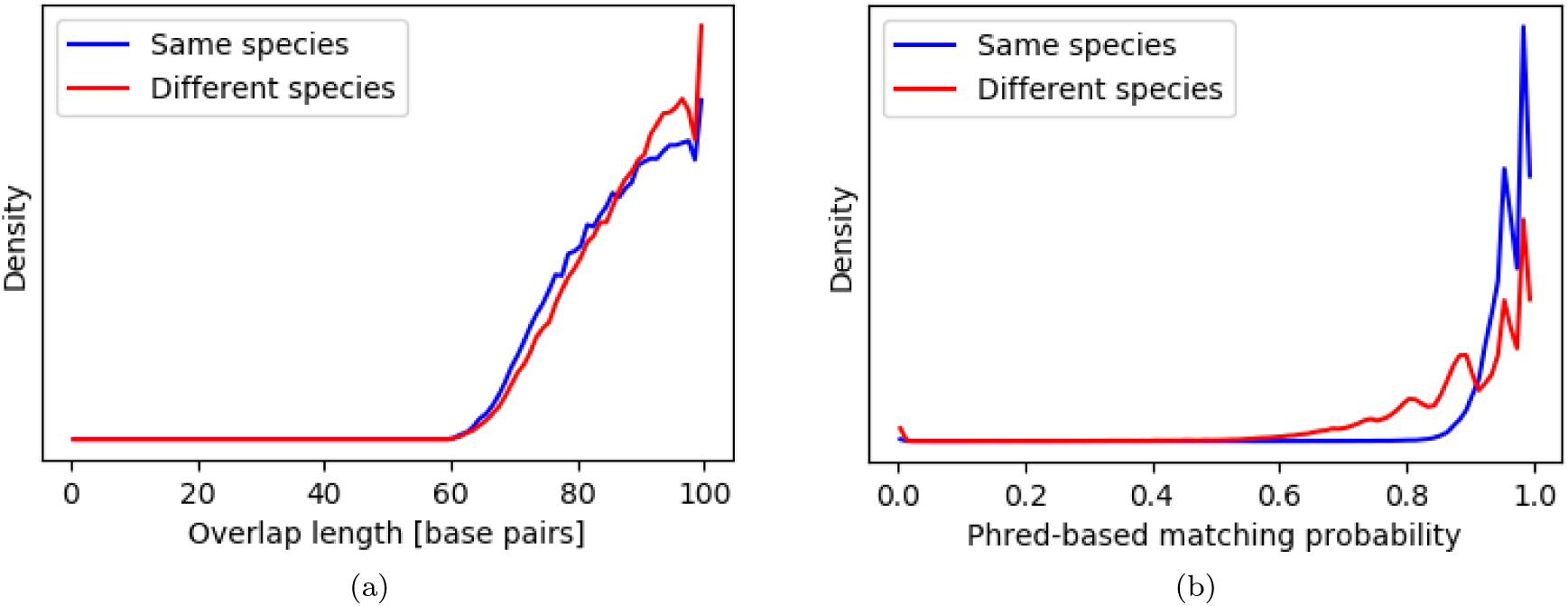
Distribution of (a) the overlap length and (b) the Phred-based matching probability for 10,000 overlaps between reads from the same species and 10,000 overlaps between reads from different species randomly selected from toy_high.

### D Clustering results: fraction of reads clustered

**Figure 13:**
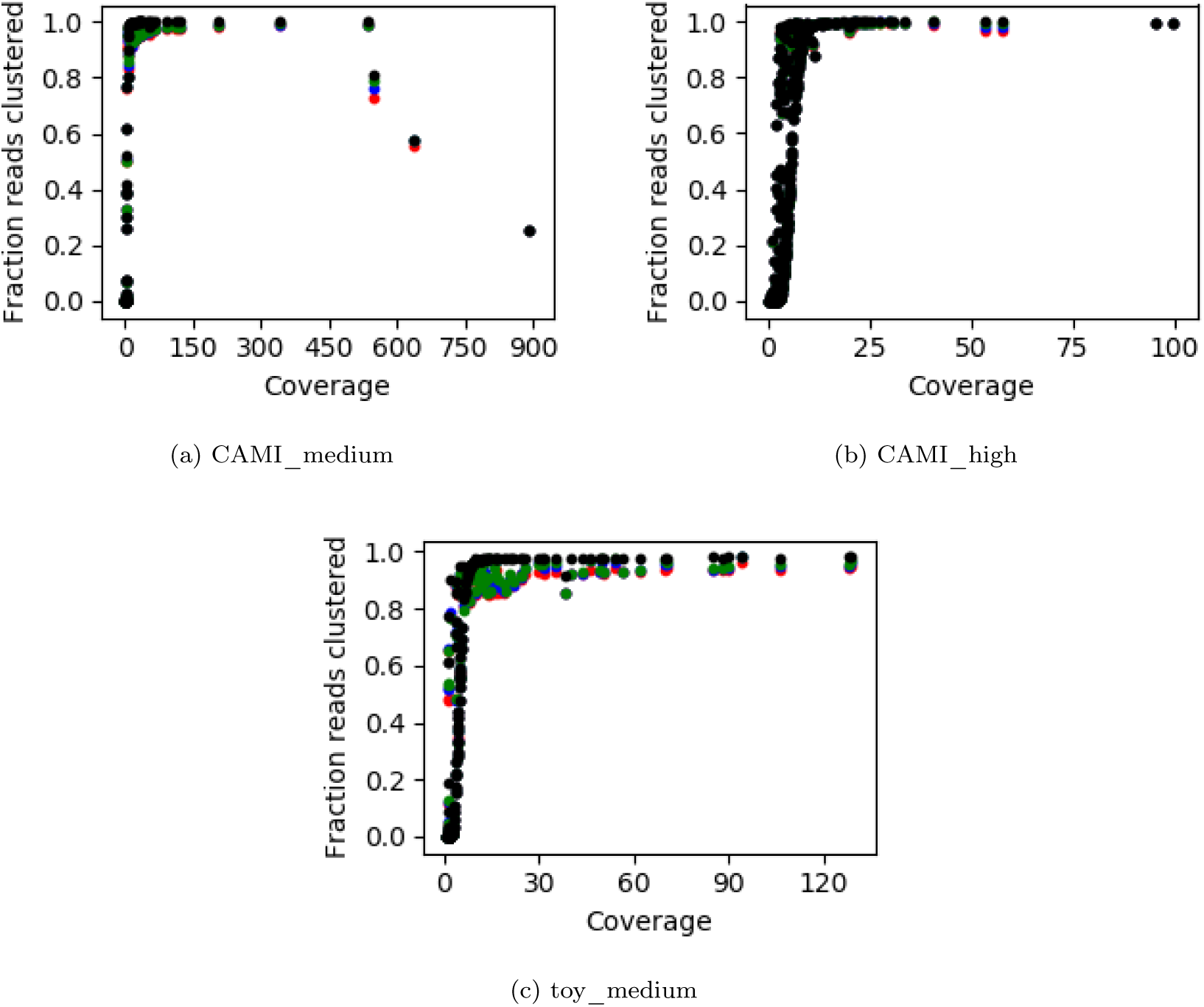
Fraction of the reads that was clustered versus coverage for (a) CAMI_medium, (b) CAMI_high, (c) toy_low and (d) toy_medium. Each dot represents a species in the dataset. Results are presented for the four maximum allowed cluster sizes: 3300 reads (red), 17000 reads (blue), 33000 reads (green) and no limit (black).

### E Clustering results: number of species per cluster

**Table 9:** Number of clusters containing the number of species indicated in the first column for CAMI_medium for the four maximum cluster sizes.

|  |  | Max cluster size (reads) |  |  |  |
| --- | --- | --- | --- | --- | --- |
|  |  | 3300 | 17000 | 33000 | Unlimited |
| No. species in cluster | 1 | 33168 | 14911 | 12167 | 3765 |
|  | 2 | 572 | 368 | 319 | 18 |
|  | 3 | 35 | 35 | 33 | 1 |
|  | 4 | 15 | 10 | 10 | 4 |
|  | 5 | 9 | 8 | 6 | 2 |
|  | 6 | 4 | 2 | 3 | 0 |
|  | 7 | 0 | 1 | 3 | 1 |
|  | 8 | 1 | 3 | 2 | 0 |
|  | 9 | 0 | 1 | 2 | 0 |
|  | 10 | 2 | 0 | 0 | 0 |
|  | 11 | 4 | 1 | 1 | 0 |
|  | 12 | 0 | 1 | 0 | 0 |
|  | 14 | 0 | 0 | 1 | 0 |
|  | 55 | 0 | 0 | 0 | 1 |
| Total |  | 33810 | 15341 | 12547 | 3792 |

**Table 10:** Number of clusters containing the number of species indicated in the first column for CAMI_high for the four maximum cluster sizes.

|  | Max cluster size (reads) |  |  |  |
| --- | --- | --- | --- | --- |
|  | 3300 | 17000 | 33000 | Unlimited |
| 1 | 203826 | 197551 | 196942 | 194155 |
| 2 | 9143 | 8223 | 8082 | 7612 |
| 3 | 1276 | 980 | 919 | 714 |
| 4 | 509 | 382 | 351 | 240 |
| 5 | 194 | 151 | 136 | 61 |
| 6 | 97 | 107 | 83 | 28 |
| 7 | 77 | 40 | 47 | 7 |
| 8 | 41 | 51 | 34 | 2 |
| 9 | 44 | 28 | 24 | 3 |
| 10 | 30 | 30 | 16 | 2 |
| 11 | 14 | 13 | 16 | 2 |
| 12 | 11 | 13 | 8 | 1 |
| 13 | 6 | 6 | 13 | 1 |
| 14 | 1 | 3 | 4 | 0 |
| 15 | 2 | 2 | 2 | 0 |
| 16 | 2 | 1 | 3 | 0 |
| 17 | 1 | 0 | 0 | 0 |
| 18 | 1 | 4 | 2 | 0 |
| 19 | 2 | 2 | 0 | 0 |
| 20 | 1 | 1 | 3 | 0 |
| 21 | 2 | 3 | 3 | 0 |
| 22 | 1 | 3 | 2 | 1 |
| 23 | 1 | 1 | 2 | 0 |
| 24 | 1 | 0 | 1 | 0 |
| 26 | 0 | 0 | 1 | 0 |
| 27 | 0 | 0 | 1 | 0 |
| 30 | 0 | 1 | 1 | 0 |
| 32 | 1 | 0 | 0 | 0 |
| 35 | 1 | 0 | 0 | 0 |
| 39 | 2 | 0 | 0 | 0 |
| 40 | 0 | 0 | 0 | 1 |
| 41 | 1 | 0 | 0 | 0 |
| 43 | 0 | 0 | 1 | 0 |
| 44 | 0 | 1 | 0 | 0 |
| 50 | 1 | 0 | 0 | 0 |
| 56 | 1 | 0 | 0 | 0 |
| 59 | 2 | 0 | 0 | 0 |
| 60 | 2 | 1 | 1 | 1 |
| 63 | 0 | 2 | 2 | 0 |
| 70 | 0 | 0 | 1 | 0 |
| 72 | 1 | 0 | 0 | 0 |
| 73 | 0 | 0 | 1 | 0 |
| 74 | 0 | 1 | 0 | 0 |
| 273 | 0 | 0 | 0 | 1 |
| Total | 215295 | 207601 | 206702 | 202832 |

**Table 11:** Number of clusters containing the number of species indicated in the first column for toy_medium for the four maximum cluster sizes.

|  |  | Max cluster size (reads) |  |  |  |
| --- | --- | --- | --- | --- | --- |
|  |  | 3300 | 17000 | 33000 | Unlimited |
| No. species in cluster | 1 | 87101 | 63553 | 59313 | 41241 |
|  | 2 | 3403 | 2322 | 2060 | 244 |
|  | 3 | 1696 | 828 | 724 | 20 |
|  | 4 | 1111 | 382 | 270 | 2 |
|  | 5 | 99 | 85 | 70 | 4 |
|  | 6 | 49 | 44 | 46 | 1 |
|  | 7 | 16 | 10 | 15 | 0 |
|  | 8 | 7 | 9 | 11 | 0 |
|  | 9 | 5 | 9 | 2 | 0 |
|  | 10 | 5 | 1 | 3 | 0 |
|  | 11 | 1 | 2 | 1 | 0 |
|  | 12 | 0 | 4 | 2 | 0 |
|  | 13 | 0 | 2 | 2 | 0 |
|  | 14 | 0 | 0 | 1 | 0 |
|  | 16 | 0 | 1 | 2 | 0 |
|  | 19 | 0 | 0 | 1 | 0 |
|  | 136 | 0 | 0 | 0 | 1 |
| Total |  | 93493 | 67252 | 62523 | 41513 |

### F Clustering results: number of clusters per species

**Figure 14:**
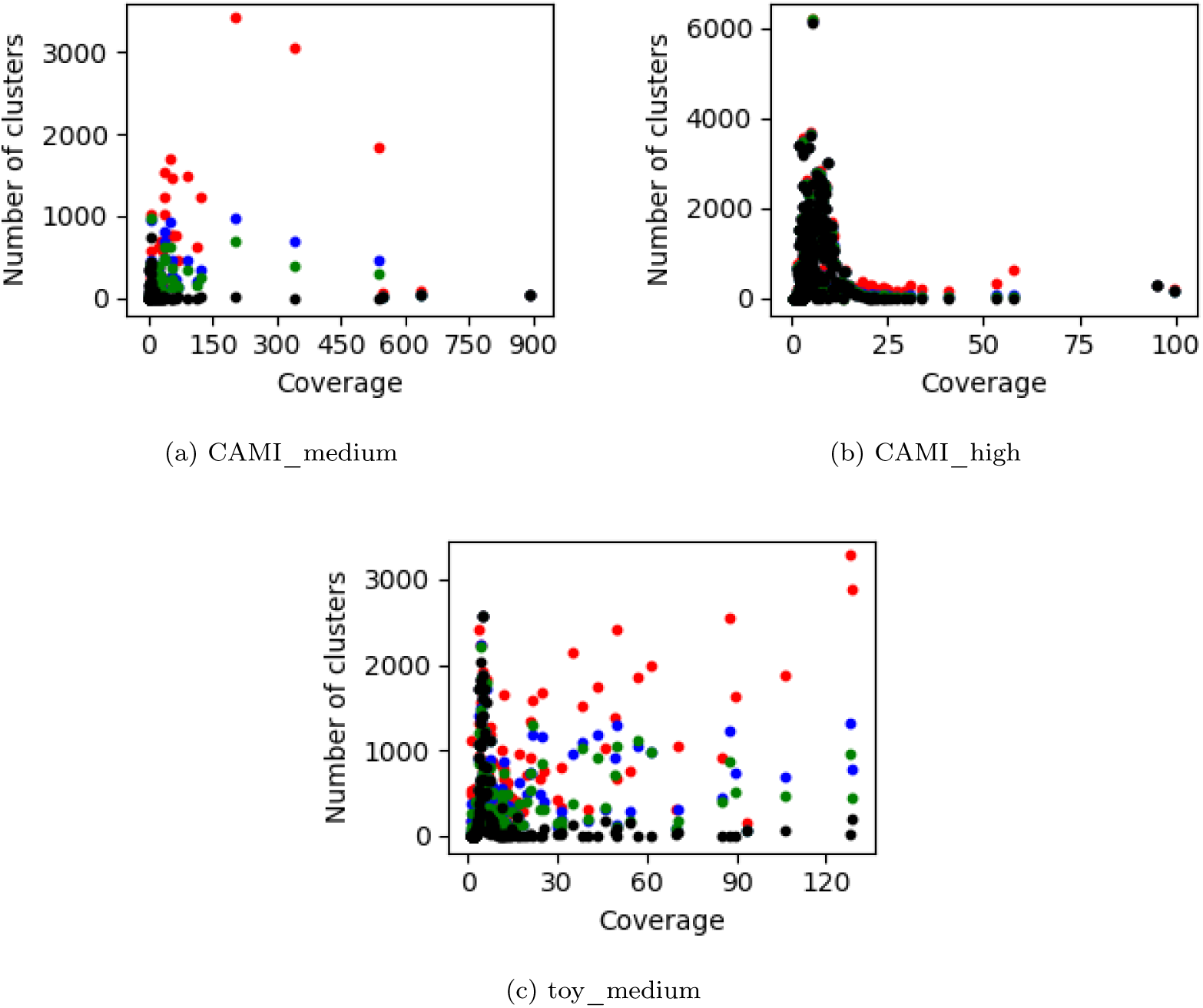
The number of clusters that contain at least one read from a species versus coverage for (a) CAMI_medium, (b) CAMI_high, (c) toy_low and (d) toy_medium. Each dot represents a species in the dataset. Results are presented for the four maximum allowed cluster sizes: 3300 reads (red), 17000 reads (blue), 33000 reads (green) and no limit (black).

